# The search behavior of terrestrial mammals

**DOI:** 10.1101/2022.12.31.521874

**Authors:** Michael J. Noonan, Ricardo Martinez-Garcia, Christen H. Fleming, Benjamin Garcia De Figueiredo, Abdullahi H. Ali, Nina Attias, Jerrold L. Belant, Dean E. Beyer, Dominique Berteaux, Laura R. Bidner, Randall Boone, Stan Boutin, Jorge Brito, Michael Brown, Andrew Carter, Armando Castellanos, Francisco X. Castellanos, Colter Chitwood, Siobhan Darlington, J. Antonio de la Torre, Jasja Dekker, Chris DePerno, Amanda Droghini, Mohammad Farhadinia, Julian Fennessy, Claudia Fichtel, Adam Ford, Ryan Gill, Jacob R. Goheen, Luiz Gustavo R. Oliveira-Santos, Mark Hebblewhite, Karen E. Hodges, Lynne A. Isbell, René Janssen, Peter Kappeler, Roland Kays, Petra Kaczensky, Matthew Kauffman, Scott LaPoint, Marcus Alan Lashley, Peter Leimgruber, Andrew Little, David W. Macdonald, Symon Masiaine, Roy T McBride, E. Patricia Medici, Katherine Mertes, Chris Moorman, Ronaldo G. Morato, Guilherme Mourão, Thomas Mueller, Eric W. Neilson, Jennifer Pastorini, Bruce D. Patterson, Javier Pereira, Tyler R. Petroelje, Katie Piecora, R. John Power, Janet Rachlow, Dustin H. Ranglack, David Roshier, Kirk Safford, Dawn M Scott, Robert Serrouya, Melissa Songer, Nucharin Songsasen, Jared Stabach, Jenna Stacy-Dawes, Morgan B. Swingen, Jeffrey Thompson, Marlee A. Tucker, Marianella Velilla, Richard W. Yarnell, Julie Young, William F. Fagan, Justin M. Calabrese

**Author notes:** This draft manuscript is distributed solely for purposes of scientific peer review. Its content is deliberative and predecisional, so it must not be disclosed or released by reviewers. Because the manuscript has not yet been approved for publication by the U.S. Geological Survey (USGS), it does not represent any official USGS finding or policy. **Materials & Correspondence:** Data and/or methods request should be sent to Michael Noonan,.

## Abstract

Animals moving through landscapes need to strike a balance between finding sufficient resources to grow and reproduce while minimizing encounters with predators ^1,2^. Because encounter rates are determined by the average distance over which directed motion persists ^1,3–5^, this trade-off should be apparent in individuals’ movement. Using GPS data from 1,396 individuals across 62 species of terrestrial mammals, we show how predators maintained directed motion ~7 times longer than for similarly-sized prey, revealing how prey species must trade off search efficiency against predator encounter rates. Individual search strategies were also modulated by resource abundance, with prey species forced to risk higher predator encounter rates when resources were scarce. These findings highlight the interplay between encounter rates and resource availability in shaping broad patterns mammalian movement strategies.

## Main

As motile organisms move through landscapes in search of food, mates, and cover, they need to strike a balance between finding sufficient resources to grow and reproduce, while also minimizing the rate at which they encounter predators^1,2^. Because of the fitness consequences of foraging success^6,7^ and predator-prey dynamics^8,9^, there should be strong selection pressure on movement strategies that maximize resource encounter rates while minimizing encounters with predators. Although it is well recognised that animals will adjust the size of their home-range areas based on resource availability^10–12^, there remains the question of how to optimally find resources within these ranges. In this context, random search models have proven influential in understanding how individual movement strategies translate to encounter rates^1,3,4,13.^ The consensus from these models is that directed (i.e., ballistic) movement leads to higher encounter rates than more tortuous (i.e., diffusive) movement^1,3,4^. This occurs because individuals that exhibit tortuous movement will tend to repeatedly search over the same areas, whereas directed motion allows individuals to search over a larger area within the same amount of time. The average distance over which ballistic motion persists, *l_v_* (in m), is thus a key determinant of encounter rates^1^ and a potent trait that individuals can optimize.

An individual’s ballistic length scale, *l_v_*, is a function of the spatial variance of their movement (*σ_p_*, in m^2^) and their positional and velocity autocorrelation timescales (*τ_p_* and *τ_v_* respectively, in sec), given by

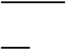

Because directed movement is the more efficient search strategy^1,4,5^, bottom-up pressure exerted by the need to encounter resources should select for more ballistic movement. Importantly, however, increasing *l_v_* will also increase the rate at which individuals encounter predators^1^. Top-down predation pressure should thus select for shorter ballistic length scales. Prey species searching for immobile vegetation must therefore optimize their movement against the opposing forces of their energetic requirements selecting for longer *l_v_*, and predation pressure selecting for shorter *l_v_*^1^. Predators also benefit from maintaining longer ballistic length scales^1^, but without the intense top-down predation pressure experienced by prey species. The combination of bottom-up and top-down regulation is thus expected to select for longer ballistic length scales in predators, versus more diffusive movement in prey species, all else being equal^1^ (Fig. 1).

**Figure 1.**
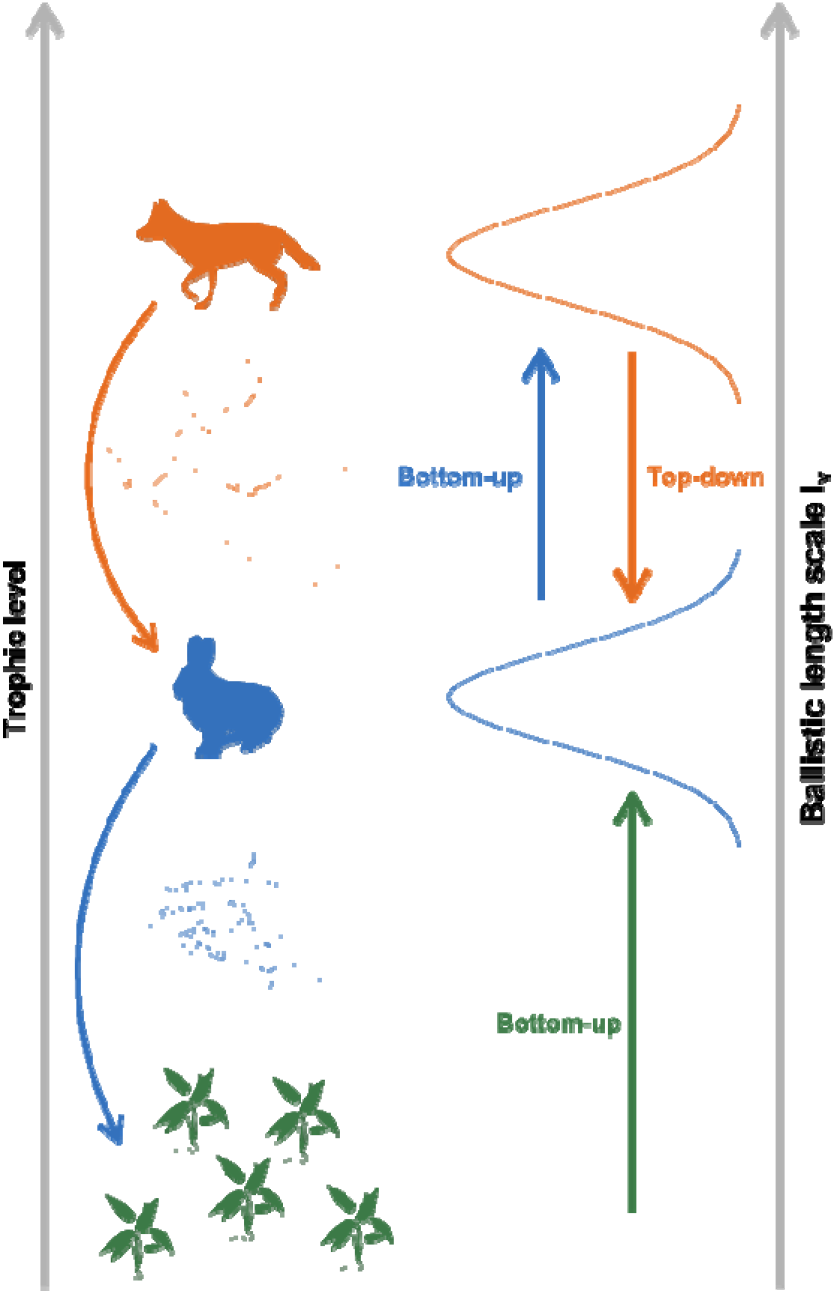
Selection pressures on predator and prey ballistic length scales. Schematic representation of bottom-up energetic requirements selecting for longer ballistic length scales in mammalian movement paths, and top-down predation pressure selecting for shorter ballistic length scales. The simulated prey movement path (blue) has a ballistic length scale of 10m, whereas the predator’s movement (orange) has a ballistic length scale of 100m.

Notably, ‘all else’ is rarely equal in ecological systems, and the relative importance of bottom-up versus top-down regulation is expected to be context specific. In resource-poor ecosystems, individuals need to spend a substantial amount of time searching for food and moving between patches^6,14^. The bottom-up driven need to find sufficient resources to survive should outweigh top-down pressure when resources are scarce. In productive environments, in contrast, bottom-up pressure on resource acquisition rates should be relaxed^12^, providing individuals with the capacity to respond more directly to top-down pressure and maintain relatively more diffusive movement. These considerations lead to the expectation of a negative relationship between *l_v_* and environmental productivity.

Although the importance of *l_v_* in governing encounter rates is suggested by theoretical models^1,5^, there has, to date, been no empirical demonstration of systematic differences in *l_v_* between comparably sized predators and prey for any taxonomic group. Here, we leverage the rapid advances in the capacity to collect^15^ and work with^16,17^ animal movement data that have enabled *l_v_* to be estimated for a broad range of species. We annotated Global Positioning System (GPS) location data on 1,396 individuals from 62 species of terrestrial mammals (Fig. 2a) with mean adult body size and trophic group (prey *n* = 41, and predator *n* = 21). We restricted our analyses to range-resident animals and used continuous-time stochastic models^16^ to estimate *l_v_* for each individual (Fig 2b). We also annotated each data point with the mean Normalized Difference Vegetation Index (NDVI), a satellite-derived measure of resource abundance^18^, to which each individual was exposed. Finally, as measures of habitat permeability we quantified the mean percent forest cover, terrain roughness, and human footprint index at each sampled location.

**Figure 2.**
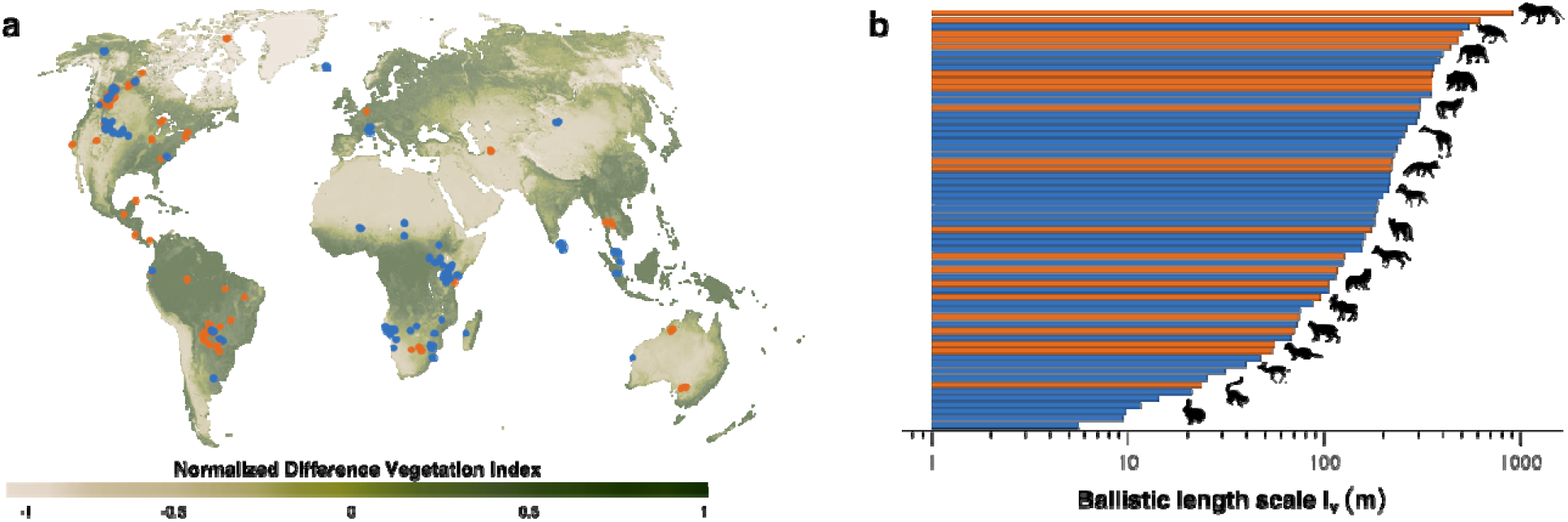
The distribution of mammalian movement data. **a**, the GPS locations of 1,396 prey (blue) and predatory (orange) mammals across 62 species are plotted on the global map of Normalized Difference Vegetation Index ranging from low (−1) to high (1) productivity; and **b** shows the median ballistic length scales, *l_v_*, for each species.

Our analyses revealed allometric scaling in ballistic length scales, with larger mammals tending to have more directed movement, all else being equal (*P* < 10^-7^; Fig. 3a). The parameters of the body-mass scaling relationships are shown in Table S2. The residuals of the body mass relationship followed theoretical predictions, with predator *l_v_* being 7.1 times longer than that of comparably sized prey species (*P* < 10^-6^; Fig. 3b). Least-squares regression also revealed a strong negative correlation between NDVI and the residuals of the allometric relationships in *l_v_* for prey species (*P* < 10^-6^; Fig. 3c). In other words, when resource abundance was ignored, predictions from the simple body-size relationships tended to under-estimate observed *l_v_* in low-productivity ecosystems, and over-estimate *l_v_* in highly productive ecosystems. That bottom-up pressures appear to outweigh top-down pressure for prey species living in resource-poor ecosystems is made all the more poignant when contrasted against known patterns in trophic structure scaling. Resource-poor environments tend to be more trophically top heavy than productive ecosystems^19^, effectively increasing the number of predators per individual prey on the landscape. Nonetheless, prey species living in resource-poor ecosystems exhibited significantly longer ballistic length scales than those in resource-rich environments, on average. Because predation rates depend on numerous factors beyond encounter rates, (e.g., capture efficiency, predator hunger levels at the time of the encounter, the presence of suitable cover, prey defenses, etc.^20^), we would expect prey species to respond more directly to bottom-up factors than predators. In line with this expectation, we found no evidence that predators adjusted their ballistic length scales as a function of NDVI (*P* = 0.26; Fig. 3C).

**Figure 3.**
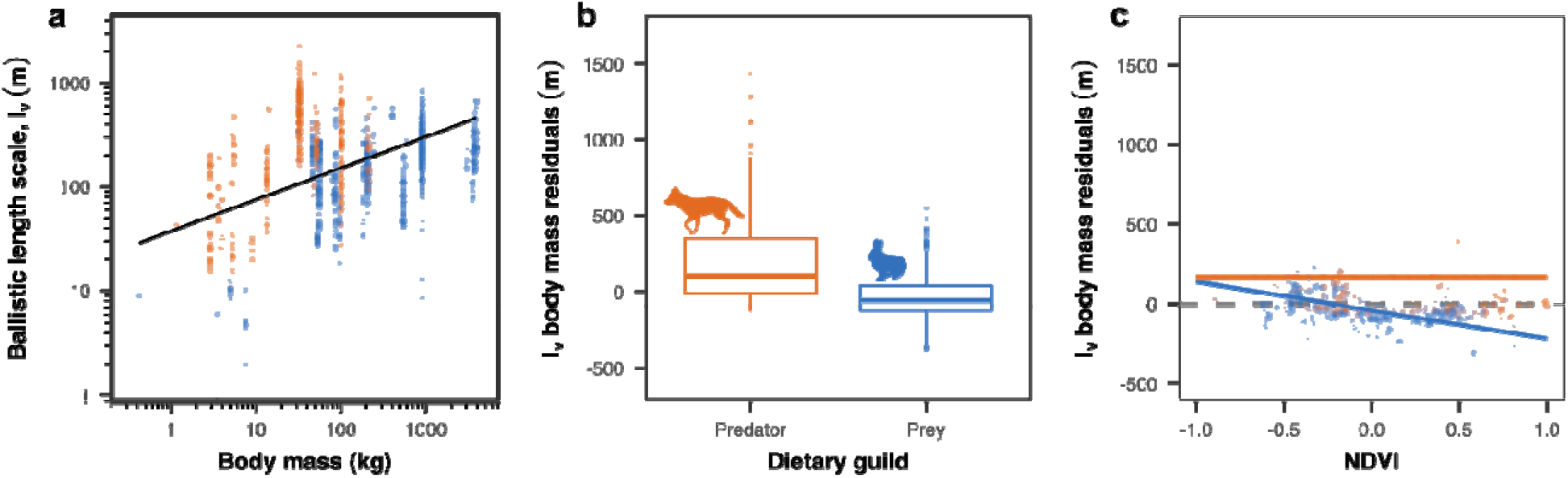
Trends in mammalian ballistic length scales. Mammalian *l_v_* scales with body size, but predators exhibit more ballistic motion than prey, and search behavior is modulated by environmental productivity. The scatterplot in **a** show the allometric scaling of the ballistic length scale, *l_v_* for prey (blue), as well as predatory (orange) mammals. The boxplots in **b** show the residuals of the body-mass scaling of *l_v_* for predators and prey. When body size is accounted for, predatory species have longer ballistic length scales on average (*P* < 10^-5^). The scatterplot in **c** show the body mass residuals as a function of the normalized difference vegetation index (NDVI). The solid lines in **c** show how prey adjusted their ballistic length scales based on NDVI (*P* < 10^-6^), whereas predators did not (*P* = 0.26).

Although predation pressure and resource abundance will influence the relative importance of search times and movement rates^6,12,14,21^, they are not the only factors that will influence animal movement. For instance, memory^22^, socially transmitted information^23^, and the ‘patchiness’ of resources^6,14^ have all been shown to influence foraging behavior. Similarly, landscape permeability can impact the capacity for individuals to maintain directed motion, with implications for foraging efficiency^24^. Indeed, we found complex, non-linear and heteroskedastic relationships between habitat structure and *l_v_* (Fig. S2). Yet, even with all of these other factors at play within the individual datasets we analyzed, our capacity to identify a clear signal for the interplay between encounter rates and resource availability in shaping mammalian movement strategies highlights just how important risk-reward trade-offs are to animals.

One question that arises from these results is ‘Why don’t predators have longer ballistic length scales?’. Without top-down regulation, there should be little preventing longer *l_v_* in predators, yet we found that predator *l_v_* was only ~7 times longer than that of comparably sized prey on average. There are two main reasons why predator ballistic length scales are not longer than observed. The first is that individuals need to perceive resources within the area they travel through, and while more directed motion might be the more efficient movement strategy, it does not necessarily imply that animals will perceive their targets. Individuals are thus constrained by their perceptual range^1,13^ and the time and effort required to find resources in the landscapes they move through^20^. Increasing *l_v_* thus only benefits predators up to a certain point, and beyond some optimal value encounter rates may actually decrease^4,5,14^. The second reason is that while many adult mammalian predators have few predators themselves, their movement can be constrained by intra-guild and conspecific encounters ^25,26^, or parasite avoidance^27^. Longer *l_v_* in predators might increase prey encounter rates, but at the expense of increasing the rates at which they encounter these negative factors.

## Conclusions

The behavioral consequences of foraging within a landscape of fear have been theorized about extensively^28,29^, yet demonstrated for only a handful of systems, such as elephant seals (*Mirounga angustirostris*), which optimize their diving behavior against predation risk and their energetic state^2^. Here, we demonstrate how the optimization of search strategies and encounter rates underpin broad patterns in the movement of terrestrial mammals spanning orders of magnitude in body size and distributed in multiple different ecosystems around the world. Prey species searching for immobile vegetation under the threat of predation must trade off the efficiency of their search strategies against the risk of encountering predators. However, ballistic length scales were negatively correlated with resource abundance, revealing how prey species are forced to risk higher predator encounter rates when resources are scarce. In contrast, the ballistic length scales of predatory species searching for mobile prey experience less top-down selection pressure, allowing them to maintain more efficient search behavior. These results highlight the interplay between encounter rates and resource availability in shaping mammalian movement strategies.

## Methods

### Empirical analyses

#### Tracking data analysis

To investigate pattern in the ballistic length scales of terrestrial mammals, we compiled openly available GPS tracking data from the online animal tracking database Movebank^30^, or from coauthors directly. Individual datasets were selected based on the criterion of range resident behavior, as evidenced by plots of the semi-variance in positions as a function of the time lag separating observations (i.e., variograms) with a clear asymptote at large time lags^16,31^. All data from migratory, or dispersing periods were excluded as their measured movement strategies would not be representative of the normal foraging dynamics we aimed to describe. The visual verification of range-residency via variogram analysis^31^ was conducted using the R package ctmm (version 1.1.0)^16^.

As noted in the main text, the average length scale over which ballistic motion is maintained (*l_v_*, in m) is a function of the spatial variance of an animal’s movement (*σ_position_*, in m^2^) and its positional and velocity autocorrelation timescales (*τ_position_* and *τ_velocity_* respectively, in sec). While can be well estimated from coarse data^32^, autocorrelation structures are only revealed when the time scale of measurement is less than or equal to the autocorrelation timescales^33^. In particular, *τ_v_* tends to be on the order of minutes to hours for medium to large terrestrial mammals^32,34^. Estimating *l_v_* for these data thus first required estimating the autocorrelation structure in each of the individual tracking datasets and identifying those for which there was sufficient information to estimate *σ_p_, τ_p_*, and *τ_v_*. To do this, we fit a series of range-resident, continuous-time movement models to the data. The fitted models included the Independent and Identically Distributed IID process, which features uncorrelated positions and velocities; the Ornstein-Uhlenbeck (OU) process, which features correlated positions but uncorrelated velocities^35^; and an OU-Foraging (OUF) process, featuring both correlated positions, and correlated velocities^31,36^. We then employed AICc based model selection to identify the best model for the data^36,37^, from which the *σ_p_, τ_p_*, and *τ_v_* parameter estimates were extracted. To fit and select the movement models, we used the R package ctmm, applying the workflow described by^16^. Because only the OUF process included information on all of the parameters required to estimate *l_v_*, we further restricted our analyses to only those individuals for which the OUF model was selected. This latter threshold was based on the sampling resolution of the GPS data, as only data of a sufficiently fine sampling resolution allow for the estimation of *l_v_*. In other words, we used AICc based model selection to identify the individual datasets with data of sufficient resolution to allow for an estimate of *l_v_*, rather than relying on an arbitrary sampling threshold. The final dataset included data from 62 species, comprising a total of 8,613,485 locations for 1,396 individuals. Finally, *l_v_* was calculated from the parameter estimates for each of these individuals as

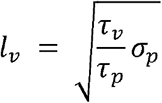

#### Covariate data

For each of the species in our dataset we compiled covariate data on that species’ mean adult mass, in kilograms, and diet taken from the EltonTraits database^38^. This dataset includes species with body masses covering five orders of magnitude (0.4 – 4000 kg). Dietary class was then used to categorize species as being either a predator or prey species. Predators were species that specialized primarily on mobile animal prey, whereas prey were herbivores and frugivores that specialized primarily on sessile vegetation. A summary of the dataset is shown in Table S1. To assess ecological factors that may have influenced *l_v_*, we also annotated each estimate with four satellite-derived habitat metrics: i) mean Normalized Difference Vegetation Index (NDVI), as a measure of local resource abundance^18^; ii) percent forest cover^39^; iii) terrain roughness^40^; and iv) machine-learning human footprint index (ml-HFI)^41^, as measures of habitat permeability. Details on the annotation process for the habitat covariate data were as follows:

1. **NDVI**. To annotate the location data with NDVI we first compiled NDVI data from the publicly accessible NASA MODIS archive. We used global NDVI rasters sampled at a 250m resolution at 16-day intervals between 2000-2022 (i.e., a total of 518 raster layers). For each individual GPS location in the dataset, we identified the point in time and space that was closest to the sampled location and extracted the NDVI value. Once each location was annotated with the appropriate NDVI value, we calculated the geometric mean value of NDVI to which each individual was exposed.
2. **Percent forest cover**. Consensus land cover data at a 1-km resolution were obtained from the EarthEnv data repository based on the methods presented in^39^. These data were available as 12 raster layers containing percentages on the prevalence of one of 12 landcover classes. From these individual raster layers we quantified the percent forest cover as the summation of five layers containing information on the prevalence of evergreen/deciduous needleleaf trees, evergreen broadleaf trees, deciduous broadleaf trees, mixed/other trees, and shrubs. Following a similar process as the NDVI annotation, we identified the point in space that was closest to each sampled GPS location and extracted the percent forest cover value and calculated the geometric mean value to which each individual was exposed.
3. **Terrain roughness**. Terrain roughness data at a 1-km resolution were obtained from the EarthEnv data repository based on the methods presented in^40^. Terrain roughness represents a measure of topographic heterogeneity, and was quantified as the largest inter-cell difference in elevation between a focal cell and its 8 surrounding cells. Here again we identified the point in space that was closest to each sampled GPS location and extracted the roughness value and calculated the geometric mean value to which each individual was exposed.
4. **ml-HFI**. The machine-learning-based human footprint index (ml-HFI)^41^ is an index of human pressure on the landscape that is derived from remotely sensed surface imagery and ranges on a scale between 0 (no human impact), and 1 (high human impact). Briefly, convolutional neural networks, are used to identify patterns of human activity from the Hansen Global Forest Change imagery version 1.7 (GFCv1.7). The raster is available at an approximately 1-km resolution. We identified the point in space that was closest to each sampled GPS location and extracted the ml-HFI value and calculated the geometric mean value to which each individual was exposed.

**Table S1.** Data summary table. Summary statistics on the GPS tracking data used in the analyses presented in the main text. Values include the species binomial, including subspecies where appropriate, the number of individuals per species (*n*), body mass, trophic group, the median spatial variance of the animals’ movement (*σ_p_*), the median positional (*τ_p_*) and velocity (*τ_v_*) autocorrelation timescales, and the median ballistic length scale (*l_v_*).

| Binomial | n | Mass (kg) | Trophic Group | $\sigma_p$ (km<sup>2</sup>) | $\tau_p$ (hrs) | $\tau_v$ (min) | $l_v$ (m) |
| --- | --- | --- | --- | --- | --- | --- | --- |
| *Acinonyx jubatus* | 1 | 46.7 | Predator | 14.882 | 190.21 | 98.05 | 357.58 |
| *Aepyceros melampus* | 20 | 52.5 | Prey | 0.229 | 33.43 | 8.05 | 31.12 |
| *Alces alces* | 56 | 541.46 | Prey | 8.56 | 706.57 | 31.62 | 76.29 |
| *Antidorcas marsupialis* | 10 | 31.5 | Prey | 66.451 | 345.81 | 42.96 | 338.58 |
| *Antilocapra americana* | 46 | 46.08 | Prey | 20.41 | 264.17 | 31.75 | 215.78 |
| *Beatragus hunteri* | 4 | 79.13 | Prey | 4.888 | 128.43 | 20.38 | 101.8 |
| *Blastocerus dichotomus* | 3 | 86.67 | Prey | 0.929 | 145 | 13.45 | 37.97 |
| *Brachylagus idahoensis* | 3 | 0.42 | Prey | 0.002 | 13.18 | 35.18 | 8.93 |
| *Canis latrans* | 49 | 13.41 | Predator | 2.981 | 17.6 | 6.18 | 115.15 |
| *Canis lupus* | 128 | 32.18 | Predator | 58.427 | 88.49 | 27.43 | 542.95 |
| *Capra ibex* | 39 | 85.17 | Prey | 4.155 | 353.57 | 25.02 | 74.6 |
| *Cerdocyon thous* | 19 | 5.24 | Predator | 0.156 | 7.43 | 1.63 | 21.62 |
| *Cervus canadensis* | 14 | 200 | Prey | 67.242 | 1636.76 | 27.93 | 167.19 |
| *Chlorocebus pygerythrus* | 10 | 4.99 | Prey | 0.024 | 11.05 | 3.35 | 9.94 |
| *Connochaetes taurinus* | 35 | 180 | Prey | 41.062 | 490.7 | 12.12 | 118.85 |
| *Cuon alpinus* | 11 | 14.17 | Predator | 5.589 | 51.68 | 33.05 | 244.06 |
| *Elephas maximus indicus* | 24 | 3500 | Prey | 50.144 | 1375.75 | 48.63 | 124.34 |
| *Elephas maximus maximus* | 51 | 3750 | Prey | 12.971 | 202.4 | 64.2 | 232.89 |
| *Elephas maximus sumatranus* | 9 | 3000 | Prey | 26.06 | 1013.66 | 51.66 | 177.53 |
| *Equus hemionus hemionus* | 18 | 240 | Prey | 1129.502 | 4715.34 | 14.21 | 234.94 |
| *Equus quagga* | 9 | 400 | Prey | 463.718 | 930.14 | 34.95 | 494.23 |
| *Eulemur rufifrons* | 3 | 2.25 | Prey | 0.029 | 7.03 | 10.83 | 24.01 |
| *Euphractus sexcinctus* | 4 | 4.78 | Prey | 0.024 | 4.85 | 1.19 | 11.23 |
| *Felis catus* | 53 | 2.88 | Predator | 2.05 | 74.06 | 8.72 | 80.7 |
| *Felis silvestris* | 3 | 5.1 | Predator | 2.634 | 30.4 | 23.67 | 108.93 |
| *Giraffa camelopardalis antiquorum* | 17 | 899.99 | Prey | 48.821 | 676.22 | 28.25 | 210.75 |
| *Giraffa camelopardalis camelopardalis* | 51 | 899.99 | Prey | 18.352 | 195.06 | 33.92 | 232.42 |
| *Giraffa camelopardalis peralta* | 17 | 899.99 | Prey | 66.868 | 253.61 | 33.64 | 346.69 |
| *Giraffa giraffa angolensis* | 72 | 899.99 | Prey | 39.839 | 303.38 | 36.98 | 255.91 |
| *Giraffa giraffa giraffa* | 31 | 899.99 | Prey | 6.082 | 118.25 | 34.9 | 183.03 |
| *Giraffa reticulata* | 35 | 899.99 | Prey | 20.083 | 191.98 | 31.17 | 211.52 |
| *Giraffa tippelskirchi* | 18 | 899.99 | Prey | 9.298 | 157.74 | 27.83 | 172.82 |
| *Hyaena brunnea* | 3 | 42.98 | Predator | 19.798 | 44.85 | 25.77 | 425.29 |
| *Lagostrophus fasciatus* | 1 | 1.7 | Prey | 0.005 | 6.1 | 34.35 | 21.17 |
| *Leopardus pardalis* | 3 | 11.9 | Predator | 0.246 | 7.66 | 9.85 | 72.59 |
| *Loxodonta africana* | 22 | 3940.03 | Prey | 242.666 | 1025.23 | 34.06 | 309.1 |
| *Lycalopex culpaeus* | 8 | 8.62 | Predator | 0.505 | 5.26 | 5.65 | 95.17 |
| *Lynx rufus* | 13 | 8.9 | Predator | 0.506 | 22.71 | 2.19 | 28.56 |
| *Madoqua guentheri* | 15 | 7.5 | Prey | 0.003 | 2.96 | 1.69 | 4.86 |
| *Neogale vison* | 5 | 1.15 | Predator | 0.548 | 127.94 | 53.65 | 44.45 |
| *Odocoileus hemionus* | 5 | 54.21 | Prey | 26.912 | 1734.41 | 23.69 | 91.76 |
| *Odocoileus virginianus* | 33 | 55.51 | Prey | 0.187 | 12.04 | 16.09 | 69.39 |
| *Oreamnos americanus* | 4 | 72.5 | Prey | 4.618 | 48.56 | 53.35 | 318.42 |
| *Oryx dammah* | 38 | 200 | Prey | 38.379 | 265.81 | 20.29 | 203.16 |
| *Ovis canadensis* | 3 | 74.64 | Prey | 10.951 | 822.88 | 47.67 | 114.35 |
| *Ovis dalli* | 66 | 55.65 | Prey | 23.112 | 524.31 | 61.06 | 204.6 |
| *Panthera leo* | 3 | 161.5 | Predator | 31.722 | 57.14 | 65.13 | 748.76 |
| *Panthera onca* | 105 | 100 | Predator | 11.749 | 126.68 | 45.69 | 254.45 |
| *Panthera pardus pardus* | 7 | 48.75 | Predator | 29.444 | 359.11 | 13.47 | 101.31 |
| *Panthera pardus saxicolor* | 6 | 52.04 | Predator | 15.643 | 185.1 | 30.3 | 221.34 |
| *Pekania pennanti* | 17 | 4 | Predator | 0.707 | 20.18 | 2.93 | 35.31 |
| *Propithecus verreauxi* | 7 | 3.48 | Prey | 0.006 | < 0.01 | < 0.01 | 6.11 |
| *Puma concolor* | 31 | 51.6 | Predator | 46.227 | 428.86 | 52.92 | 264.8 |
| *Rangifer tarandus* | 19 | 86.03 | Prey | 55.734 | 1521.59 | 62.11 | 182.18 |
| *Rangifer tarandus tarandus* | 8 | 86.03 | Prey | 285.467 | 1749.55 | 72.86 | 402.56 |
| *Sus scrofa* | 23 | 96.12 | Prey | 0.234 | 7.53 | 2.18 | 43.85 |
| *Syncerus caffer* | 6 | 580 | Prey | 23.035 | 255.58 | 29.75 | 240.2 |
| *Tapirus terrestris* | 41 | 207.5 | Prey | 0.367 | 9.11 | 21.41 | 117.39 |
| *Tremarctos ornatus* | 1 | 140 | Prey | 15.035 | 258.17 | 35.41 | 185.38 |
| *Ursus arctos horribilis* | 18 | 212.5 | Predator | 30.3 | 249.65 | 40.92 | 253.13 |
| *Vulpes lagopus* | 18 | 3.58 | Predator | 1.289 | 4.42 | 0.81 | 64.09 |
| *Vulpes vulpes* | 20 | 5.48 | Predator | 1.39 | 5.89 | 13.04 | 199.24 |

#### Assessing trends in l_v_

The resulting dataset of ballistic length scales was then analyzed to test for differences in *l_v_* between predators and prey, and for any effects of NDVI or forest cover on *l_v_*. Because *l_v_* was correlated with body size (Fig. 3A), we controlled for mass by regressing *l_v_* against body size on a log_10_-log_10_ scale using generalized least-squares fitting with Gaussian distributed errors. Due to phylogenetic autocorrelation^42^, closely related species may exhibit similarities in movement due to common descent. Accordingly, we did not treat species data records as independent, but rather corrected for this inertia by adjusting the variance-covariance matrix in our regression model based on the phylogenetic relationships using the `R` package `nlme`^43^. Phylogenetic relationships between the mammalian species in our dataset were obtained from the VertLife repository^44^. We sampled 1,000 trees and estimated the consensus tree using the `R` package `phytools`, version 1.0-3^45^. Several of the species and subspecies included in our analyses were not included in the VertLife repository. These were manually inserted into the phylogeny using the most recent estimates of divergence times from the closest related species found in the VertLife repository. This included the Persian leopard (*Panthera pardus saxicolor*), with divergence time of 0.297 Ma from the African leopard (*P. pardus pardus*)^46^, the Sumatran elephant (*Elephas maximus sumatranus*), with divergence time of 190,000 years from the Indian elephant (*E. maximus indicus*)^47^, the Sri Lankan elephant (*E. maximus maximus*), with divergence time of 43,000 years from the Indian elephant^47^, elk (*Cervus canadensis*), with divergence time of 20 million years from *Rangifer tarandus*^48^, and Norwegian reindeer (*R. tarandus tarandus*) with a divergence time from *R. tarandus* of 115,000 years^49^. Divergence times for all of the species of *Giraffa* were taken from^50^. The species *G. camelopardalis* was assigned a divergence time from *G. giraffa* of 0.37 Ma. *G. tippelskirchi* was assigned a divergence time from *G. giraffa* of 0.23 Ma, and *G. reticulata* a divergence time from *G. camelopardalis* of 0.26 Ma. The subspecies *G. giraffa angolensis* and *G. giraffa giraffa* had a divergence time of 0.04 Ma. *G. camelopardalis antiquorum* was assigned a divergence time from *G. camelopardalis peralta* of 0.15 Ma and *G. camelopardalis camelopardalis* from *G. camelopardalis peralta* of 0.12 Ma. The resulting phylogenetic tree is shown in figure S1.

The final step in our analyses was to determine whether individual deviations from the allometric relationship in *l_v_* could be described by trophic group, ecosystem productivity, or habitat permeability. We regressed the residuals of the phylogenetically controlled allometric model described above against trophic group, NDVI, percent forest cover, terrain roughness, and ml-HFI using generalized least-squares fitting with Gaussian distributed errors. The relative support for these models was then assessed by comparing the AICc values of the fitted models against intercept only models. We chose to work with the residuals as it allowed for like-to-like comparisons across all of the individuals in our dataset and clearer visualisations of the partial effects, while still correcting for differences in body sizes and phylogenetic inertia. The parameters of the body-mass scaling relationships in *l_v_* are shown in Table S2. Results of the trophic group and NDVI regressions are presented in the main text and the model selection results are presented in Table S3. The R scripts used to produce the results presented in this work are openly available on GitHub at https://github.com/NoonanM/BallisticMotion.

**Table S2.** Observed scaling relations of ballistic length scales in terrestrial mammals. All three of the scaling relations had the general form *l_v_* = *β_0_* mass*^α^*.

| | $\beta_0$ | 95% CI | $\alpha$ | 95% CI |
| --- | --- | --- | --- | --- |
| **All taxa** | 4.67 | 1.34 — 15.72 | 0.30 | 0.19 — 0.41 |
| **Prey** | 0.84 | 0.22 — 3.22 | 0.42 | 0.30 — 0.53 |
| **Predators** | 1.26 | 0.19 — 8.56 | 0.51 | 0.32 — 0.71 |

**Figure S1.**
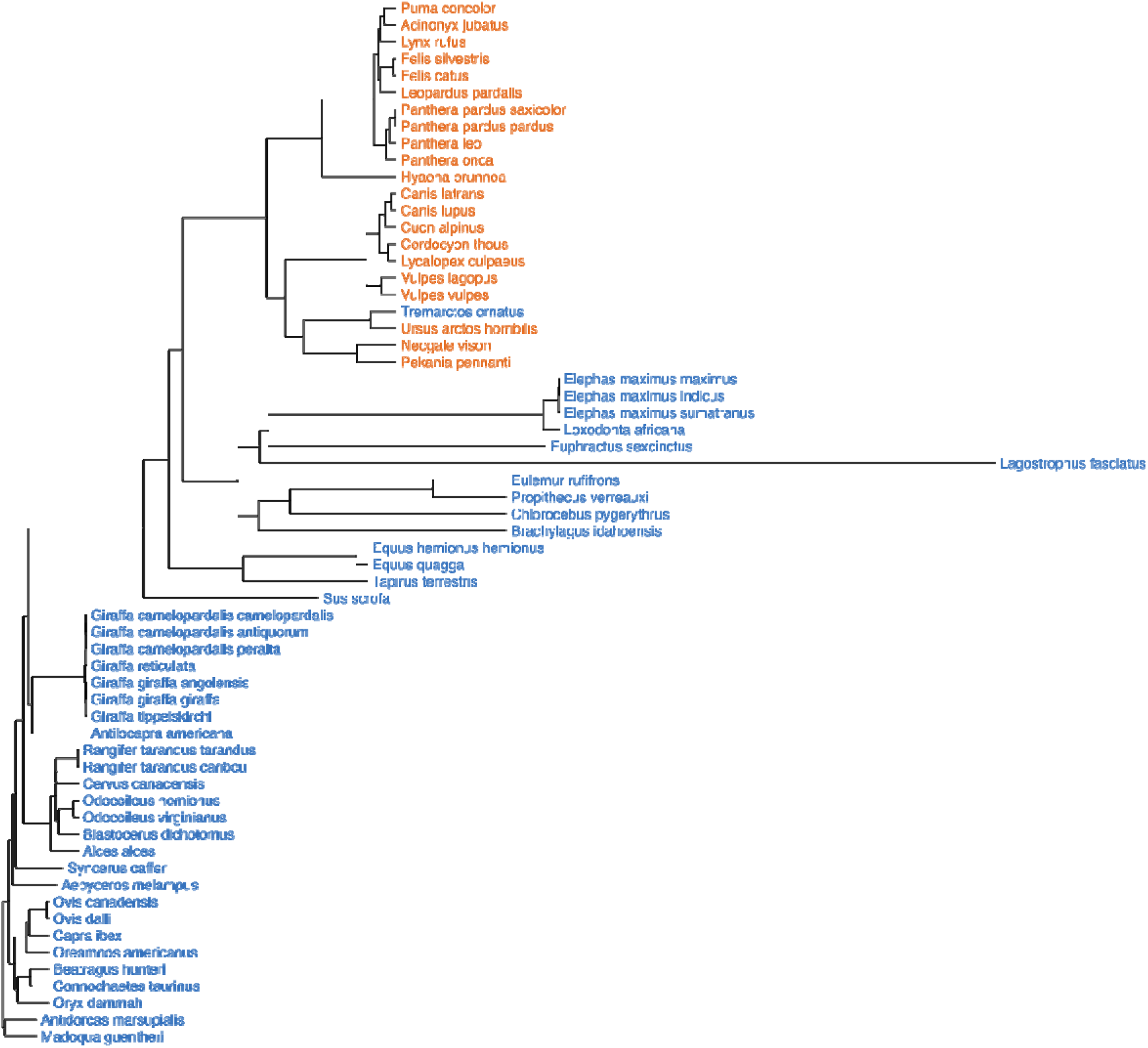
Phylogenetic relationships of the Mammalian species analysed in the main text with species labels coloured by trophic guild (prey in blue, predators in orange).

**Table S3.**
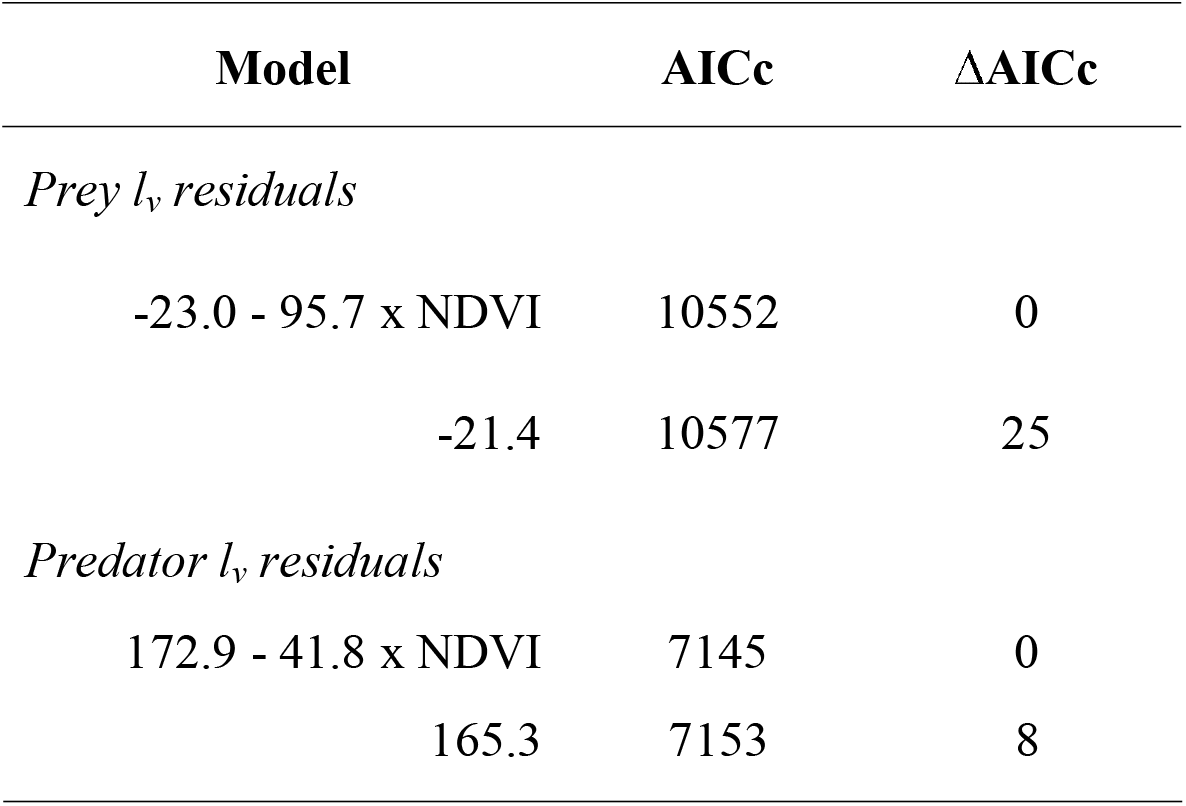
Table showing the model selection results for the effect of NDVI on the residuals of the body mass relationship in ballistic motion length scales, *l_v_*, for predators and prey.

When corrected for body size, mammalian ballistic length scales, *l_v_*, show mixed responses to measures of habitat permeability. We found no relationship between the *l_v_* body mass residuals and percent tree cover within each individual’s habitat for prey (*P* = 0.06) nor predators (*P* = 0.92), but both trophic groups exhibited exponentially variable heteroskedasticity with changing forest cover (Fig. S2a, table S4). We also found no relationship between the *l_v_* body mass residuals and terrain roughness for prey (*P* = 0.06), but there was a positive relationship for predators (*P* = 0.001). Here again both trophic groups exhibited exponentially variable heteroskedasticity with changing terrain roughness (Fig. S2b, table S5). Finally, both predators and prey living in human modified landscapes had shorter ballistic length scales than those living in more natural environments, but these relationships were non-linear and heteroskedastic (Fig. S2c, table S6). Although this latter result is somewhat surprising given some empirical studies’ findings that the structure present in natural ecosystems can impact the capacity for individuals to maintain directed motion^24^, it is in line with the reductions in mammalian movement observed in human modified environments generally^21^.

**Figure S2.**
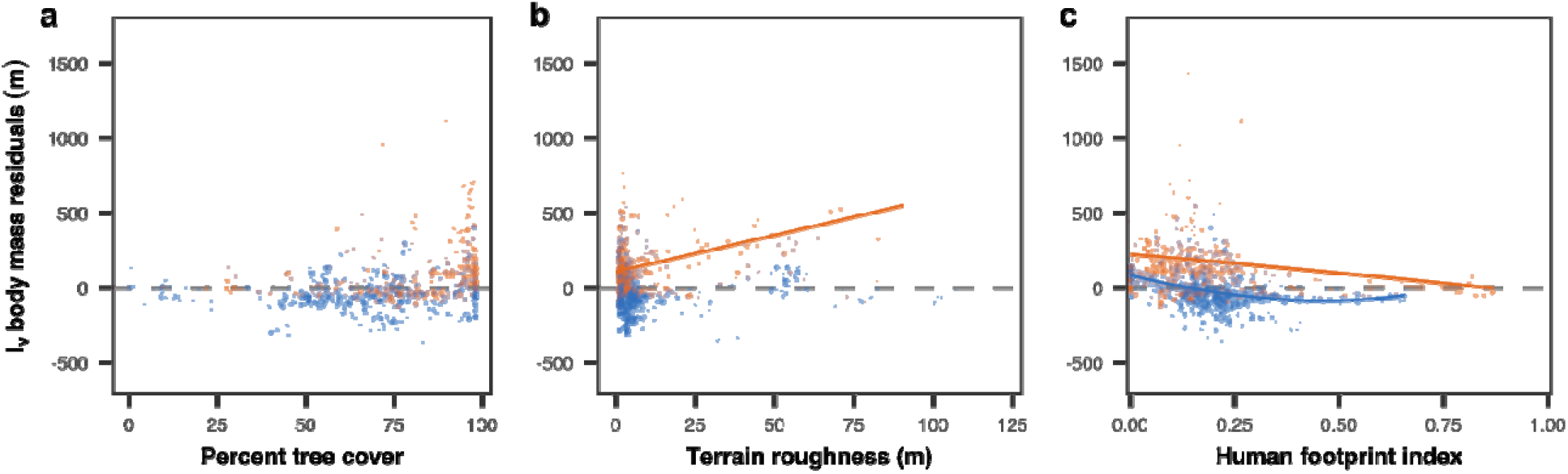
Relationship between mammalian ballistic length scales and habitat permeability. When corrected for body size, mammalian ballistic length scales, *l_v_*, show mixed responses to measures of habitat permeability. The scatterplot in **a** shows the residuals of the allometric scaling of the ballistic length scale, *l_v_* for prey (blue), as well as predatory (orange) mammals as a function of the percent tree cover. In **b** the residuals of the body mass scaling of *l_v_* for predators and prey are shown as a function of terrain roughness. The scatterplot in **c** depicts the body mass residuals as a function of the human footprint index, a satellite derived measure of human disturbance ranging on **a** scale between 0 and 1. Each point is an individual (*n* = 1,396) representing 62 species. The solid lines depict the selected regression models and the shaded areas the 95% confidence intervals around the mean trend. Trends are not shown for cases where the intercept-only model was selected.

**Table S4.** Table showing the model selection results for the effect of percent forest cover on the residuals of the body mass relationship in ballistic motion length scales, *l_v_*, for predators and prey.

| Model | AICc | $\Delta AICc$ |
| --- | --- | --- |
| *Prey $l_v$ residuals* |  |  |
| $-14.5 + \varepsilon \sim N(0, \sigma^2 \times e^{-0.004 \times \text{forest cover}})$ | 10574 | 0 |
| $-53.0 + 0.5 \times \text{forest cover} + \varepsilon \sim N(0, \sigma^2 \times e^{-0.004 \times \text{forest cover}})$ | 10574 | 0 |
| $-52.7 + 0.5 \times \text{forest cover} + \varepsilon \sim N(0, \sigma^2)$ | 10577 | 3 |
| $-21.4 + \varepsilon \sim N(0, \sigma^2)$ | 10577 | 3 |
| *Predator $l_v$ residuals* |  |  |
| $158.7 + \varepsilon \sim N(0, \sigma^2 \times e^{0.014 \times \text{forest cover}})$ | 7145 | 0 |
| $163.7 - 0.1 \times \text{forest cover} + \varepsilon \sim N(0, \sigma^2 \times e^{0.014 \times \text{forest cover}})$ | 7145 | 0 |
| $156.6 + 0.1 \times \text{forest cover} + \varepsilon \sim N(0, \sigma^2)$ | 7154 | 9 |
| $165.3 + \varepsilon \sim N(0, \sigma^2)$ | 7154 | 9 |

**Table S5.** Table showing the model selection results for the effect of terrain roughness on the residuals of the body mass relationship in ballistic motion length scales, *l_v_*, for predators and prey.

| Model | AICc | $\Delta\text{AICc}$ |
| --- | --- | --- |
| *Prey $l_v$ residuals* |  |  |
| $-37.4 + \varepsilon \sim N(0, \sigma^2 \times e^{-0.02 \times \text{roughness}})$ | 10504 | 0 |
| $-11.8 - 0.6 \times \text{roughness} + \varepsilon \sim N(0, \sigma^2 \times e^{-0.02 \times \text{roughness}})$ | 10505 | 1 |
| $-5.1 - 1.1 \times \text{roughness} + \varepsilon \sim N(0, \sigma^2)$ | 10563 | 59 |
| $-21.4 + \varepsilon \sim N(0, \sigma^2)$ | 10577 | 73 |
| *Predator $l_v$ residuals* |  |  |
| $106.4 + 4.9 \times \text{roughness} + \varepsilon \sim N(0, \sigma^2 \times e^{0.02 \times \text{roughness}})$ | 7127 | 0 |
| $130.9 + \varepsilon \sim N(0, \sigma^2 \times e^{0.02 \times \text{roughness}})$ | 7138 | 11 |
| $102.2 + 4.3 \times \text{roughness} + \varepsilon \sim N(0, \sigma^2)$ | 7141 | 14 |
| $165.3 + \varepsilon \sim N(0, \sigma^2)$ | 7154 | 27 |

**Table S6.**
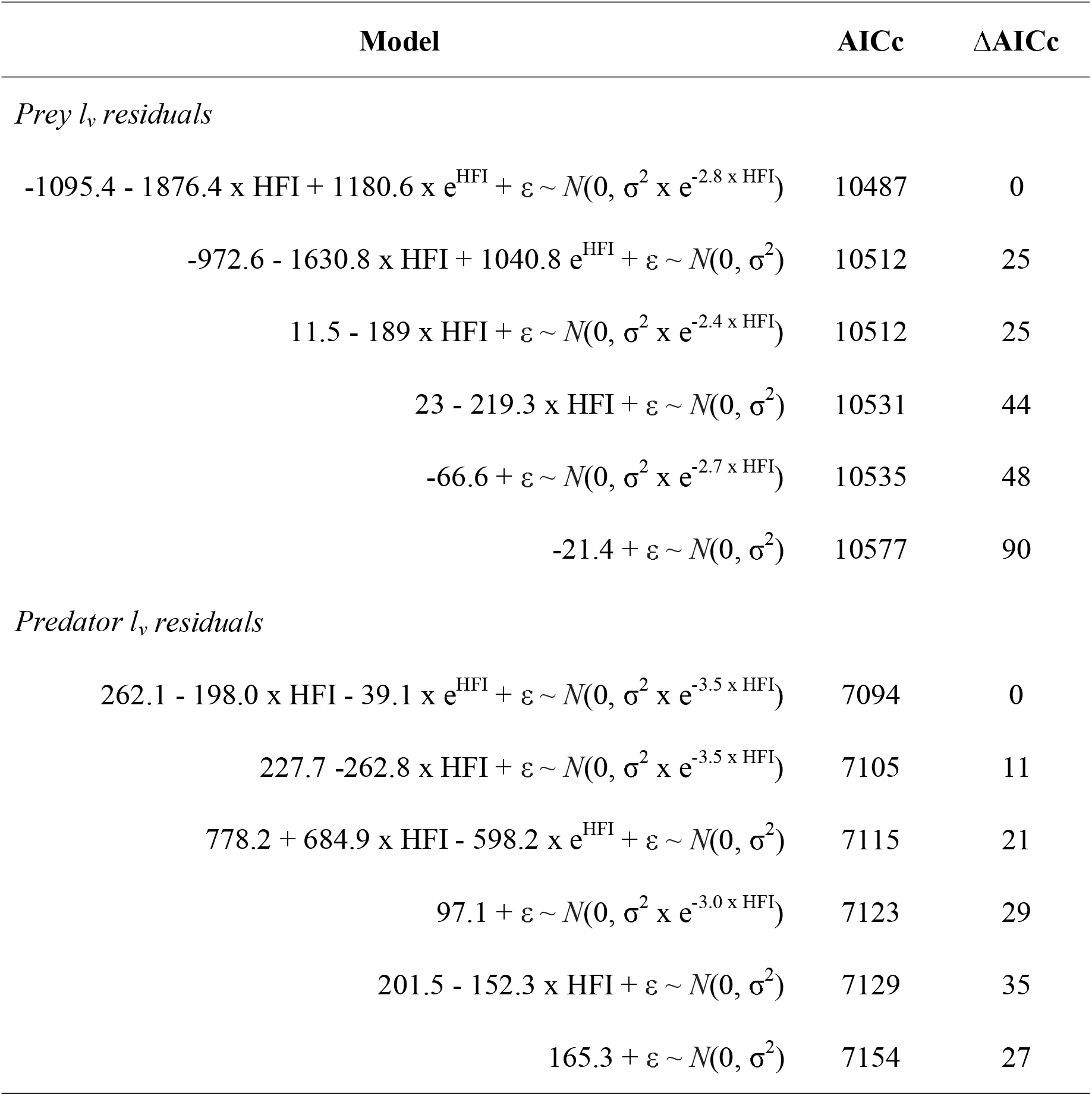
Table showing the model selection results for the effect of the human footprint index on the residuals of the body mass relationship in ballistic motion length scales, *l_v_*, for predators and prey.

## Acknowledgments

The full list of funding sources for this work is provided in Appendix S1. Any use of trade, firm, or product names is for descriptive purposes only and does not imply endorsement by the U.S. Government.

## Author contributions

MJN and JMC conceived the manuscript; MJN conducted the analyses. MJN, JMC, RMG, CHF, and WHF wrote the first manuscript draft. Co-authors contributed data and assisted with writing the final version of the manuscript.

## Data and materials availability

The data reported in this paper and the R scripts used to carry out this study are openly available on GitHub at https://github.com/NoonanM/BallisticMotion.

## Supplementary Materials

Figs. S1 to S2

Tables S1 to S6

Appendix S1: Extended List of Acknowledgements and Funders

Data S1: Data used to generate the results presented in the main text.

## Appendix S1: Extended List of Acknowledgements and Funders

MJN was supported by an NSERC Discovery Grant RGPIN-2021-02758. This work was partially funded by the Center of Advanced Systems Understanding (CASUS) which is financed by Germany’s Federal Ministry of Education and Research (BMBF) and by the Saxon Ministry for Science, Culture and Tourism (SMWK) with tax funds on the basis of the budget approved by the Saxon State Parliament. CHF, WFF, and JMC were supported by NSF IIBR 1915347. R.M.-G. was supported by FAPESP through grant no. ICTP-SAIFR 2016/01343-7; Programa Jovens Pesquisadores em Centros Emergentes through grant nos. 2019/24433-0 and 2019/05523-8, and the Simons Foundation through grant number 284558FY19; BGF was supported by FAPESP through grant 2019/26736-0; R.M.-G and B.G.F were supported by Instituto Serrapilheira through grant no. Serra-1911-31200. The procurement of the collars in Ecuador was funded by the International Climate Initiative of the German Federal Ministry for the Environment, Nature Conservation, and Nuclear Safety with support of KfW Development Bank (BMZ project no. 2098 10 987). MSF was supported by a research fellowship from the University of Oxford and the Iranian Department of Environment approved permits for the work conducted (93/16270). Financial support was provided by the People’s Trust for Endangered Species (PTES), Zoologische Gesellschaft für Arten und Populationsschutz (ZGAP), Iranian Cheetah Society, IdeaWild and Association Francaise des Parcs Zoologiques (AFdPZ) to the leopard collaring project in Iran. RGM was supported by FAPESP grants 2013/10029-6 and 2014/24921-0. RK was supported by USA NSF grants 2206783, 1914928, 1754656 and NASA Ecological Forecasting Program Grant 80NSSC21K1182. DB was supported by Canada Foundation for Innovation (grant no. 38881), Natural Sciences and Engineering Research Council of Canada (grants no. 509948-2018, RGPIN-2019-0592, RGPNS-2019-305531), and Network of Centers of Excellence of Canada ArcticNet. LAI thanks the US National Science Foundation (grant nos. BCS 99-03949 and BCS 1266389), L.S.B. Leakey Foundation, and Committee on Research, University of California, Davis, for financial support, the Kenya Government (NACOSTI permit No. P/15/5820/4650) for research permission, and the Kenya Wildlife Service for local affiliation. The Dutch Wildcat data was obtained in a project for ARK Nature with funding of the province of Limburg. JJT, RTM, and MV were supported by PROCIENCIA project 14□INV□208 from the Consejo Nacional de Ciencia y Tecnología of Paraguay with the permission of the Ministerio del Medio Ambiente y Desarrollo Sostenible of Paraguay under research permits 124/2017 and 155/2018. Funding for research on introduced predators was provided by supporters of Australian Wildlife Conservancy with additional support from the Australian Government’s National Environmental Science Program through the Threatened Species Recovery Hub. Movement data for scimitar-horned oryx were obtained through a reintroduction project in Chad: this project is a joint initiative of the Government of Chad and the Environment Agency - Abu Dhabi, implemented in Chad by SaharaConservation in partnership with the Ministry for the Environment, Fisheries and Sustainable Development, with technical support from the Zoological Society of London, Fossil Rim Wildlife Center, Saint Louis Zoo, and other key partners. Movement data for the four giraffe species was funded by the Giraffe Conservation Foundation and its supporters, and the collaborative efforts of government, academic and NGO partners. Funding for this project was provided by the U.S. Department of Defense through the Wildlife Branch at Fort Bragg Military Installation and Fisheries, Wildlife, and Conservation Biology Program at North Carolina State University. Funding was provided by the National Science Foundation (DEB-1146166, to J.L. Rachlow) Funding was provided by the Austrian Science Fund (P18624). Funding was provided by the Utah Division of Wildlife Resources.

